# Haplotype-Resolved Assembly for Synthetic Long Reads Using a Trio-Binning Strategy

**DOI:** 10.1101/2020.06.01.126995

**Authors:** Mengyang Xu, Lidong Guo, Xiao Du, Lei Li, Li Deng, Ou Wang, Ming Ni, Huanming Yang, Xun Xu, Xin Liu, Jie Huang, Guangyi Fan

## Abstract

The accuracy and completeness of genome haplotyping are crucial for characterizing the relationship between human disease susceptibility and genetic variations, especially for the heterozygous variations. However, most of current variations are unphased genotypes, and the construction of long-range haplotypes remains challenging. We introduced a *de novo* haplotype-resolved assembly tool, HAST that exports two haplotypes of a diploid species for synthetic long reads with trio binning. It generates parental distinguishing *k*-mer libraries, partitions reads from the offspring according to the unique markers, and individually assembles them to resolve the haplotyping problem. Based on the stLFR co-barcoding data of an Asian as well as his parental massive parallel sequencing data, we utilized HAST to recover both haplotypes with a scaffold N50 of >11 Mb and an assembly accuracy of 99.99995% (Q63). The complete and accurate employment of long-range haplotyping information provided sub-chromosome level phase blocks (N50 ∼13 Mb) with 99.6% precision and 94.1% recall on average. We suggest that the accurate and efficient approach accomplishes the regeneration of the haplotype chromosomes with trio binning, thus promoting the determination of haplotype phase, the heterosis of crossbreeding, and the formation of autopolyploid and allopolyploid.

## Introduction

The whole-chromosome genetic haplotyping is of great importance for the analyses of the correlation between genotypes and phenotypes^1-3^. Haplotyping denotes the combination of allelic variants along each homologous chromosome. The detections of small structural variants or single-nucleotide variants require local genomic information, while those of large-scale structural variants such as chromosomal rearrangements need additional long-range molecular information.

Inbreeding^4^ or haploid-cloning^5^ provides a simply way to avoid the problem of haplotyping at its source. However, it is technically impractical for most species. Another strategy refers to the whole-chromosome isolation, for instance, chromosome sorting^6^, which is not applicable to large sample size due to its time-consuming and labor-intensive procedure. Haplotyping information can also be straightforwardly obtained from the sequencing of fragmented or whole DNA molecules. This approach requires long-range information to reconstruct the haplotypes from the biological or physical markers or overlapping relations of the segmented homologous chromosomes, including synthetic long reads (SLR, such as MGI stLFR co-barcoding reads and 10X genomics linked reads)^7-10^, single-molecule long reads (Pacific Biosciences (PacBio) or Oxford Nanopore (ONT) long reads)^11, 12^, Hi-C^13, 14^ and optical mapping (Bionano)^15^. In practice the long-range information is employed to recover diploid or polyploid genomes with read-mapping-based, or *de novo* assembly-based methods.

Currently, to attribute the successive genotypes to an arbitrary haplotype, sequencing reads are generally mapped to the reference genomes and then the alignment information recovers the heterozygous genotypes at each polymorphic site^16, 17^. Nevertheless, the read-mapping-based methods are hampered by the lack of perfect reference sequence and the inability to detect large structural variations. Long reads^18, 19^ or SLR^7, 8, 20^ can extend the phase blocks to ∼1Mb, but it is still complex to span regions of low heterozygous density or gaps in the reference. Assembly-based methods usually generate longer phase blocks across homozygous regions of the chromosome due to the assembly procedure of reads, and therefore can detect large indels or structural variants. A number of graph-based approaches, including FALCON-Unzip^11^, reserve the variant-induced “bubble” structures in the assembly graph, and align long reads back against the graph to arbitrary cluster one arm of each bubble and the traversal to extent the phase blocks to several Mb level. However, these approaches cannot avoid the possible switch errors between adjacent phase blocks and mis-assemblies on account of repeated allelic sites^21^. Supernova^9^ also groups bubbles in the assembly graph which share the same barcodes to obtain long continuous phase blocks. Recent efforts to address the problem include the elimination of switch errors with additional long-range information such as Hi-C and Bionano optical mapping^22-24^.

Trio-binning-assembly-based strategy significantly simplifies the haplotyping problem by classifying the offspring reads into haplotype-specific groups prior to assembly, according to the unshared heterozygous markers from the parents^12, 25, 26^. It resolves the phasing problem induced by complex allelic variations in the assembly-based approaches at the source. This approach has been recently extended to gamete binning by single-cell sequencing^27^. Although it is limited by the additional requirement of pedigree information, it has been reported that trio binning produces the long-read assemblies with improved haplotyping continuity and accuracy^12, 28^. In addition, the independent assemblies in parallel overcome the significant memory and CPU consumption, especially for complex polyploid genomes.

The utilization of SLR, such as stLFR^7, 10^, has obtained a series of successes in haplotyping^10, 29-33^, structural variation detection^33-36^ and *de novo* genome assembly^37-43^. Its achievements are established on the efficient clustering of short reads with long-range barcode information. However, there is no protocol available for SLR data to resolve the haplotyping using trio-binning stratagem.

In this work, we developed a tool, <u>H</u>aplotype-resolved <u>A</u>ssembly for <u>S</u>ynthetic long reads using a <u>T</u>rio-binning strategy (HAST, https://github.com/BGI-Qingdao/HAST), that employs the haplotype-specific *k*-mers from parents to partition the SLR long fragments, and individually *de novo* assembles them to accurately construct the haplotypes with the long-range information from SLR barcodes. The results of haplotype reconstruction are comparable to PacBio or ONT assemblies by TrioCanu^12^.

## Results

### Haploid partition of long fragment reads

We demonstrated that HAST successfully partitioned SLR long fragment reads (LFR) for the diploid genome using trio binning. The workflow utilizes the information of parental distinct heterozygous variants to facilitate a completely haplotype-resolved assembly for the diploid genome, in the following three steps (**Fig. 1**): a) generation of parental unshared *k*-mers using massive parallel sequencing (MPS) reads and SLR sequencing of the offspring; b) classification of offspring’s long fragments based on the parental unshared *k*-mer sets; and c) haplotype-resolved assemblies for both parent-inherited chromosomes.

**Figure 1.**
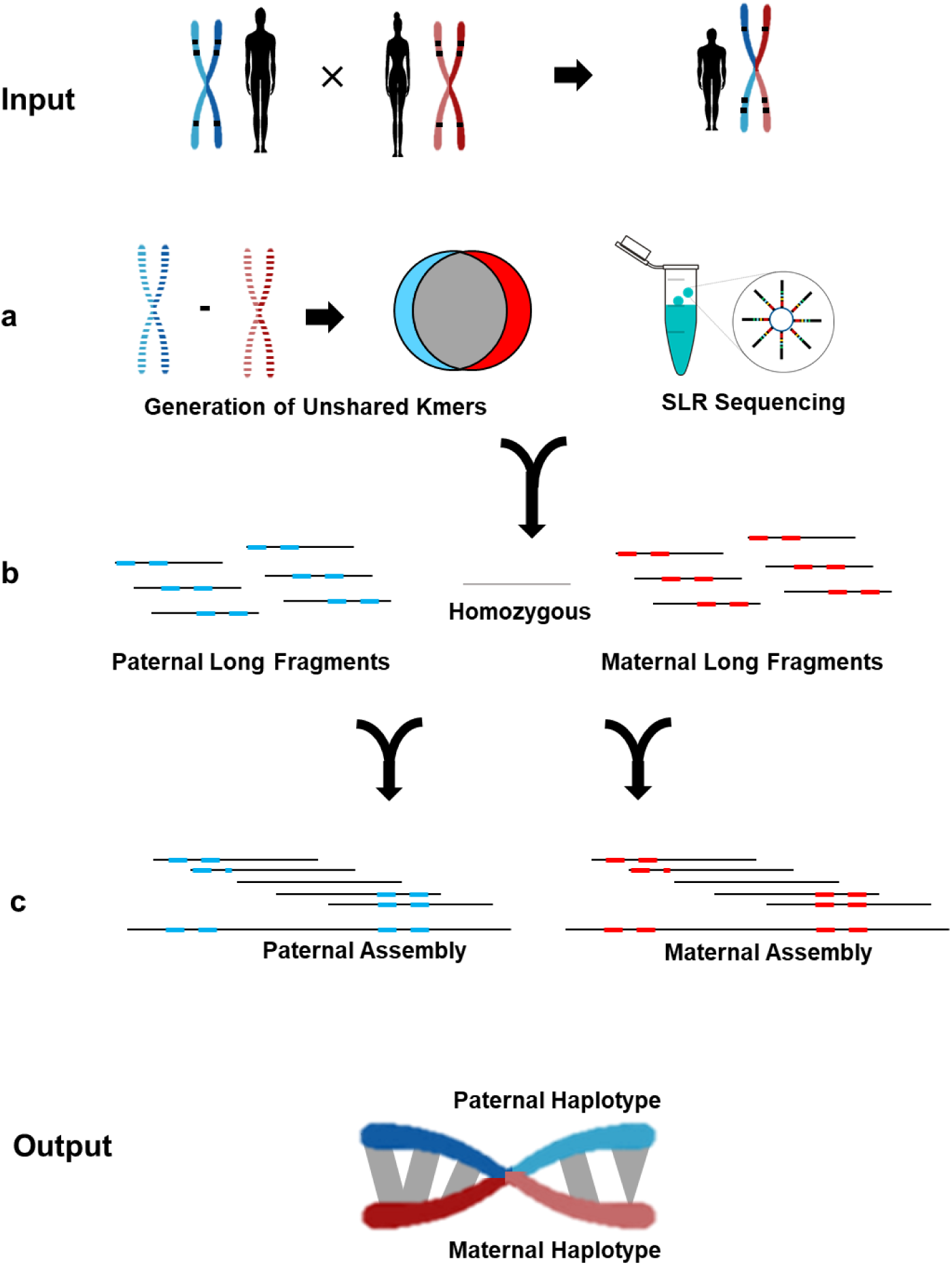
A schematic of HAST workflow. **a**) Generation of parent unshared *k*-mers, and SLR sequencing of the offspring. **b**) Classification of offspring’s long fragments based on the parental unshared *k*-mer sets. **c**) Haplotype-resolved assemblies for both parent-inherited chromosomes. The output includes two complete haplotypes.

The parental MPS read sets with ∼30× coverage for each were used to generate their unique markers (unshared *k*-mers) to classify the offspring’s SLR data to the corresponding haplotypes. The accurate selection of haplotype-specific *k*-mers remarkably depends on the sequencing error rate, and influences the precision of partitioning. In this study, we literately applied *k*=21, *k*=19 and *k*=31 to the SLR data identification to enhance the efficiency. The 21-mers that occurred less than 9 times or more than 58 were filtered out according to the *k*-mer distribution (**Supplementary Fig. 1**), resulting in ∼40 M unique markers for maternal and ∼60 M for paternal, which influenced the capability and accuracy to capture phased long fragments (**Supplementary Table 1**). In view of the limit of read length (100 bp), only ∼1.4% short reads had at least one unshared *k*-mers (**Supplementary Table 2**). However, the stLFR barcodes clustered short reads and extended the haplotyping information to the whole long fragment. There were totally 155,196,643 barcodes in the offspring’s clean reads, of which 18.7% were uniquely assigned into haplotypes. To investigate the relatively low assignment rate, we calculated the assignment rate of different stLFR groups as a function of the number of read pairs in the LFR. We observed that the ratio of LFR’s with no parental-specific *k*-mers were exponentially reducing from 93% down to 0% with the increasing fragment length, while the number of each assigned LFR groups was evenly growing (**Fig. 2a**), suggesting that the distinguishable LFR’s had relatively more read pairs and these unassigned LFR’s were short fragments with no heterozygous sites. Those with equal numbers of both parental markers seem ambiguous, but the ratio remains ∼0%. As a result, when we masked the short fragments (i.e. <20 read pairs per barcode), there were totally 31.0% barcodes (42.0% read pairs) and 32.7% barcodes (44.1% read pairs) classified as inherited from the offspring’s father or mother, respectively, indicating the feasibility and high efficiency of trio binning strategy using stLFR data.

**Figure 2.**
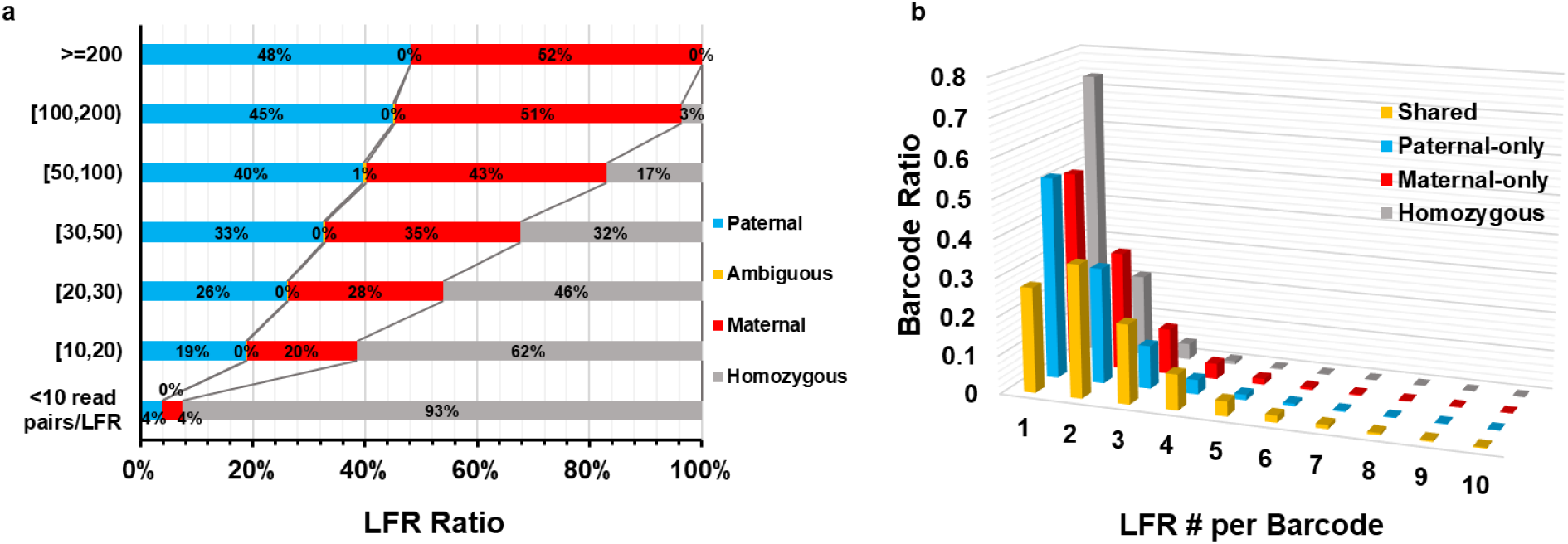
The statistics for classified LFR’s. **a**) The compositions in long fragments at different lengths. A large number of short LFR’s (<10 read pairs per LFR) are attributed to the homozygous, which have no haplotype-specific markers. As the length increases, the major portion changes to the parental-specific groups. Those with equal numbers of both parental-specific *k*-mers are classified as ambiguous (∼0%), which seem like false partitions. **b**) The LFR collision for parental-inherited, shared, and homozygous partitions. The y-axis represents the ratio of total barcodes for each dataset, and x-axis represents the number of LFR’s per barcode. Unlike others, the shared LFR’s reveal a peak at 2, indicating a higher possibility of collision. Shared: have both types of parental-specific *k*-mers; Paternal-only: only have paternal-specific *k*-mers; Maternal-only: only have maternal-specific *k*-mers; Homozygous: have no parental-specific *k*-mers.

### Collision rates of different assigned stLFR groups

To gain a full understanding of long-fragment property for the partitioning, each group was independently mapped against the human reference genome (hg19). The combination of each haploid and homozygous long fragments covered 83.0% and 87.7% reference genome, respectively, with at least 30× sequencing coverage (**Supplementary Table 3**). The mapping results also allows the evaluation of long fragment collision, which is raised by the opportunity that more than one DNA nanoballs are loaded in the same co-barcoding vessel. The stLFR collision rate refers to the average number of SLR’s from different regions of genome (reference assembly) sharing the same barcode. Paternal, maternal, shared and homozygous groups represented a 1.58, 1.54, 2.12 and 1.19 collision rate, respectively (**Supplementary Table 4**). The distribution of long fragment count per barcode showed that the majority of paternal, maternal and homozygous barcodes had only one long fragment, while the shared barcodes exhibited a peak at two (**Fig. 2b**). Those SLR’s having both haploid markers were also classified based on the relative ratios, in order to tolerate sequencing errors. Although those reads tend to introduce defective long-range information, the data proportion is not statistically significant thanks to the high base-calling accuracy of short-read-based SLR techniques. Moreover, the spurious information could be avoided by the assembly process considering the fact that the barcode collision randomly occurs among the long fragments and thus those fragments with duplicated barcodes hardly exist in the same region of the genome.

### Complete reconstruction of individual haplotypes

We individually assembled the classified stLFR data from an Asian (HJ), to reconstruct each haplotype, and compared to the direct pseudo-haplotyping assemblies according to the assembly graph structure instead of trio binning. The entire assemblies generated by HAST were in phase and the origin of each sequence was unambiguous, while the pseudo-haplotypes arbitrarily combined one arm of the “bubble” structures in the graph and had no origin information. Although it was complicated to maintain the phasing information based on the sparse heterozygous variant sites for human in consideration of the low heterozygosity (∼0.1%), HAST clustered the fragments based on the long-range barcode information and successfully recovered the haploid chromosomes. The complexity would decrease for genomes with higher heterozygosity.

The maternal-inherited heterozygous long fragments along with homozygous generated a complete assembly of 3.0 Gb with the scaffold N50 of 18.3 Mb, while the paternal assembly of 3.0 Gb provided a scaffold N50 of 11.4 Mb (**Supplementary Table 7**). With aligning to the human reference genome, the two haplotypes represented a scaffold NGA50>1 Mb and genome fraction>94.39%, which indicated the high assembly accuracy and completeness. The difference in the assembly quality between paternal and maternal could possibly arose from the complication of chromosome Y assembly. In contrast, each pseudo-haplotype assembly without trio binning produced a comparable scaffold N50 (16.7 Mb) but a shorter contig N50, which seemed no difference in the quality between two assemblies. The more complex assembly process introduced a high probability of assembly errors. Meanwhile, the *k*-mer-based estimation provided the short-range *k*-mer-level accuracy (QV), and the reappearance capacity of reliable *k*-mers from input offspring reads (completeness). HAST revealed a slightly higher average QV values (Q63 vs. Q61), and a similar completeness (∼97%) (**Table 1**).

**Table 1.** Haplotype-resolved assembly quality statistics for the *H. sapiens* HJ. The metrics in continuity were reported by QUAST. The assembly quality and haplotyping were evaluated by Merqury, including *k*-mer-based quality value (QV), assembly completeness, haplotyping precision and recall. The consensus *QV* = −10*log*_10_*E* where E refers to the single-base-level assembly errors. The haplotyping estimates were given by the reliable haplotype-specific *k*-mers. BUSCO v3 was run with vertebrata_odb9 database, where Comp.: Complete, Dup.: Duplicated, Frag.: Fragmented, Mis.: Missing.

| Assembly | Continuity |  |  |  | Quality |  | Haplotyping |  |  | BUSCO (%) |  |  |  |
| --- | --- | --- | --- | --- | --- | --- | --- | --- | --- | --- | --- | --- | --- |
|  | Scaffold | Scaffold | Contig | Contig | QV | Completeness | Precision | Recall | F1-score | Comp. | Dup. | Frag. | Mis. |
|  | Max | N50 | Max | N50 |  |  |  |  |  |  |  |  |  |
|  | (Mb) | (Mb) | (Mb) | (Mb) | (Phred) | (%) | (%) | (%) |  |  |  |  |  |
| **HAST** |  |  |  |  |  |  |  |  |  |  |  |  |  |
| Paternal | 59.7 | 11.4 | 0.61 | 0.06 | 61.83 | 96.61 | 99.34 | 94.02 | 96.61 | 91.1 | 1.4 | 5.3 | 3.6 |
| Maternal | 70.9 | 18.3 | 0.51 | 0.06 | 65.13 | 96.83 | 99.89 | 94.22 | 96.97 | 91.7 | 1.3 | 4.9 | 3.4 |
| Both | 70.9 | 14.9 | 0.61 | 0.06 | 63.18 | 99.08 | 99.62 | 94.13 | 96.80 | 98.2 | 83.6 | 1.1 | 0.7 |
| **Supernova** |  |  |  |  |  |  |  |  |  |  |  |  |  |
| Pseudohap1 | 105.1 | 16.7 | 0.34 | 0.04 | 60.62 | 97.10 | 52.90 | 64.91 | 58.29 | 87.2 | 1.4 | 8.3 | 4.5 |
| Pseudohap2 | 105.1 | 16.7 | 0.34 | 0.04 | 60.46 | 97.10 | 39.74 | 54.05 | 45.80 | 87.0 | 1.4 | 8.2 | 4.8 |
| Both | 105.1 | 16.7 | 0.34 | 0.04 | 60.54 | 98.23 | 46.32 | 84.11 | 59.74 | 88.1 | 85.0 | 7.6 | 4.3 |
| **TrioCanu** |  |  |  |  |  |  |  |  |  |  |  |  |  |
| Paternal | - | - | 5.7 | 0.73 | 37.39 | 91.76 | 99.90 | 96.12 | 97.97 | 88.8 | 1.4 | 6.1 | 5.1 |
| Maternal | - | - | 10.5 | 1.5 | 38.40 | 96.74 | 99.91 | 92.78 | 96.21 | 92.5 | 1.4 | 4.2 | 3.3 |
| Both | - | - | 10.5 | 1.1 | 37.88 | 99.24 | 99.91 | 94.57 | 97.17 | 97.8 | 87.2 | 1.4 | 0.8 |

### Haplotyping validation of the entire assemblies

As a sanity check, we applied Merqury^44^, a reference-free phasing assessment to all the assemblies with reliable parental and offspring’s *k*-mers. This is a novel but accurate method for the haplotyping evaluation considering that the alignment against the human reference assembly provides no phasing information and currently there have been no assemblies available for HJ’s parents. By counting the number of the expected parental-specific *k*-mers that existed in the offspring’s diploid assemblies, it demonstrated that the haplotypes recovered 94.02% of the paternal heterozygous sites, and 94.22% of the maternal (**Table 1**). This was significantly higher than the two pseudo-haplotypes without trio binning, which was only 64.91% and 54.05%.

The phasing error is related to the unexpected parental-specific *k*-mers. The precision value of the paternal and maternal assemblies was 99.34% and 99.89%, respectively. In contrast, it was only 52.90% and 39.74% for pseudo-haplotypes in spite of high base accuracy (>Q60) of the assemblies. Moreover, the HAST haplotypes contained fewer fragmented or missing BUSCO genes compared to the pseudo-haplotypes, and totally achieved 98.2% complete genes with lower duplication rate. It suggested that there were more allelic variations improperly assembled in the pseudo-haplotypes.

The diploid assembly spectra according to the haplotyped-resolved markers provided useful visualized information of haplotyping. In the copy-number plots (**Fig. 3a,b**) as a function of *k*-mer multiplicity in the offspring’s reads, the first and small 1-copy (heterozygous) peak (red) corresponded to heterozygous sites, the second and large 2-copy (homozygous) peak (blue) corresponded to homozygous sequence or haplotype-specific duplications, and higher copies (green, purple, orange) corresponded to repeats. Those *k*-mers that only occurred in the reads (grey) possibly came from the sequencing errors or indicated the missing genome region that assemblies could not achieve. The two approaches had comparable assembly completeness. Besides, there was a considerably small bar at zero multiplicity corresponding to the assembly *k*-mers absent from the reads, which agreed with the >Q60 assembly single-base level consensus. Besides, the haplotype-resolved assembly spectra and separated the contributions with offspring’s *k*-mer multiplicity. The *k*-mers that occur exclusively in one of the haplotypes represent the assembled haplotype-specific sites. The paternal assembly (blue) by HAST had an approximately identical number of *k*-mers in the heterozygous peak compared to the maternal assembly (red) (**Fig. 3d**). In contrast, the arbitrary combination of “heterozygous sites” by Supernova also made equal proportions of the peak for each pseudo-haplotype, but the total number of classified allelic variations was lower by 37% (**Fig. 3e**). The fraction of shared *k*-mers (orange) in the peak corresponds to the heterozygous variants shared by both parents, given that the human heterozygosity is only ∼0.1%. Moreover, the hap-mer blob plots described the continuity, accuracy and completeness of phased scaffolds. A number of paternal-specific *k*-mers in the scaffolds proved the accurate phasing of the HAST paternal assembly with almost no maternal markers, while the maternal assembly showed almost no paternal markers regardless of scaffold length (**Fig. 2g**). On the contrary, numerous scaffolds in each pseudo-haplotype shared both paternal and maternal-specific *k*-mers except short sequences (**Fig. 2h**). This indicated that pseudo-haplotypes could not reconstruct the true homologous chromosomes without trio binning, consistent with the haplotyping precision and recall (**Table 1**).

**Figure 3.**
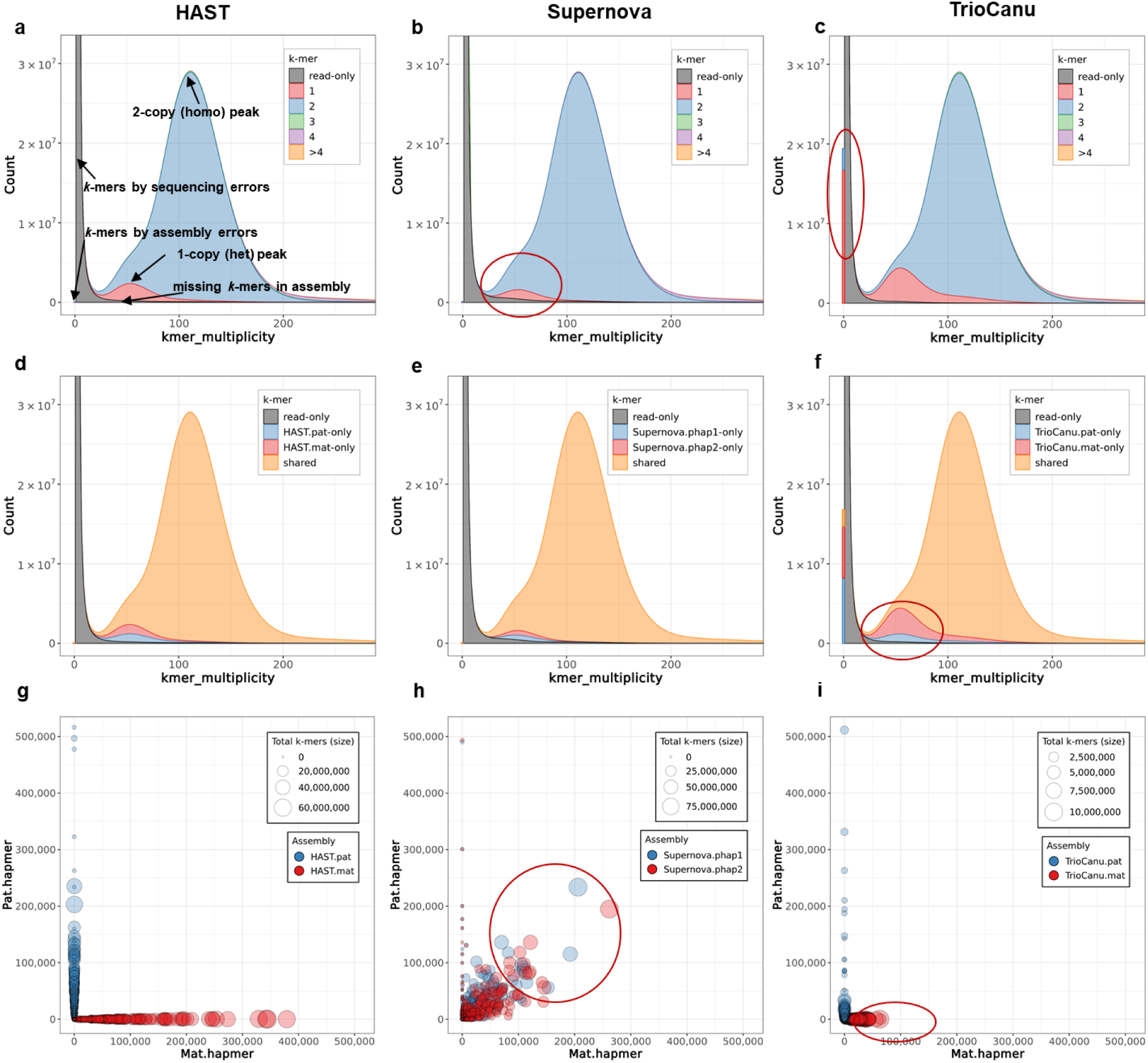
Haplotype spectra for the diploid human genome. **a-c**) The stacked copy-number spectra of *k*-mer multiplicity for evaluating assembly errors and heterozygous completeness. **d-f**) The stacked haplotyping assembly spectra of *k*-mer multiplicity for evaluating haplotyping completeness and balance. **g-i**) Hap-mer plots for evaluating haplotyping continuity and accuracy, where the”hap-mer” refers to the haplotype-specific *k*-mer for each haplotype of the genome^44^. The blob size is proportional to the scaffold/contig size, and the position is plotted based on the number of contained paternal (x-axis) and maternal (y-axis) hap-mers. (**a, d, g**) are generated by HAST-assembled stLFR data, (**b, e, h**) by directly Supernova-assembled stLFR data, and (**c, f, i**) by TrioCanu-assembled PacBio CCS data.

The continuity and accuracy of phase block are guaranteed by the reservation of long-range phasing information. However, switch errors of the assembly split the scaffold/contig and reduce the block size in practice. With the advantage of parental information, the phase block size N50 by HAST were up to 10.0 Mb and 15.7 Mb for paternal and maternal haplotypes, respectively, which were close to the scaffold N50 (**Table 1** and **Table 2**). However, the pseudo-haplotypes revealed a 10-fold decrease in the continuity from scaffolds to phase blocks, limited by the indistinguishable long-range information. By comparison, HAST represented a ∼10-fold larger phase block size relative to pseudo-haplotypes, and the longest phase block was up to the chromosome level (66.5 Mb; **Table 2**). Meanwhile, the HAST haplotypes covered almost the entire genome with low switch error rate. Overall, the HAST assemblies were more structurally accurate than the direct pseudo-haplotypes.

**Table 2.**
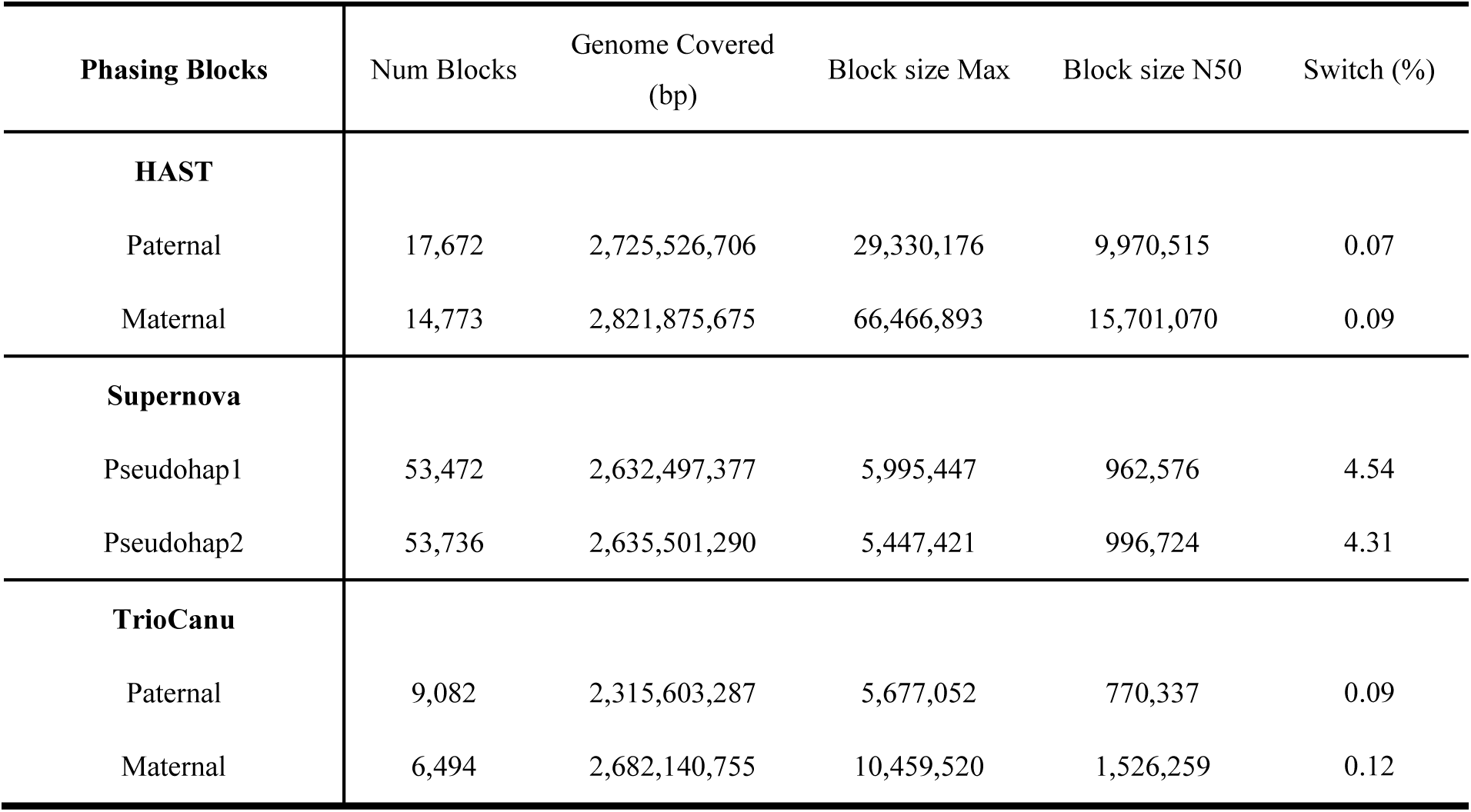
Statistics for phase blocks and switch error rates. A phase block refers to the region in the sequence where at least two same type of paternal or maternal-specific *k*-mers occur. Two consecutive different parental-specific *k*-mers within a certain range are marked as a switch error and split the sequence into two phase blocks. We allowed at most 100 switches within 20 kbp range to calculate the phase block.

### Comparisons with PacBio HiFi CCS reads and haplotypes

To verify the haplotyping effect, we also used PacBio circular consensus sequence (CCS) long reads from the same diploid sample, and examined the phased structure variations (SV) by HAST and Supernova assemblies. There were no reference assemblies for both parents currently available. Therefore, the relatively long and accurate CCS reads after consensus were ideal to inspect the haplotypes and the associated phased SV. We individually mapped the paternal or maternal CCS reads against each haplotype or pseudo-haplotype, and investigated several regions corresponding to known SV from standard calls. Each group of partitioned CCS reads was well aligned with the corresponding HAST’s haplotype (**Fig. 4**), indicating a good concordance of phased sequences. In contrast, there were gaps of approximately 460 bp (**Fig. 4a**), or overlaps of 315 bp (**Fig. 4b**) in the mapped CCS reads for one of the pseudo-haplotypes by Supernova, while the other had a great alignment. Because of the close resemblance in the structure shared by two pseudo-haplotypes and the capability of CCS reads to represent the true chromosomal configuration, the “gaps” reflected a phased deletion in the paternal-inherited chromosome 4, which was accurately captured by HAST. Meanwhile, the “overlaps” illustrated a maternal-specific insertion, which was missing in both Supernova’s assemblies.

**Figure 4.**
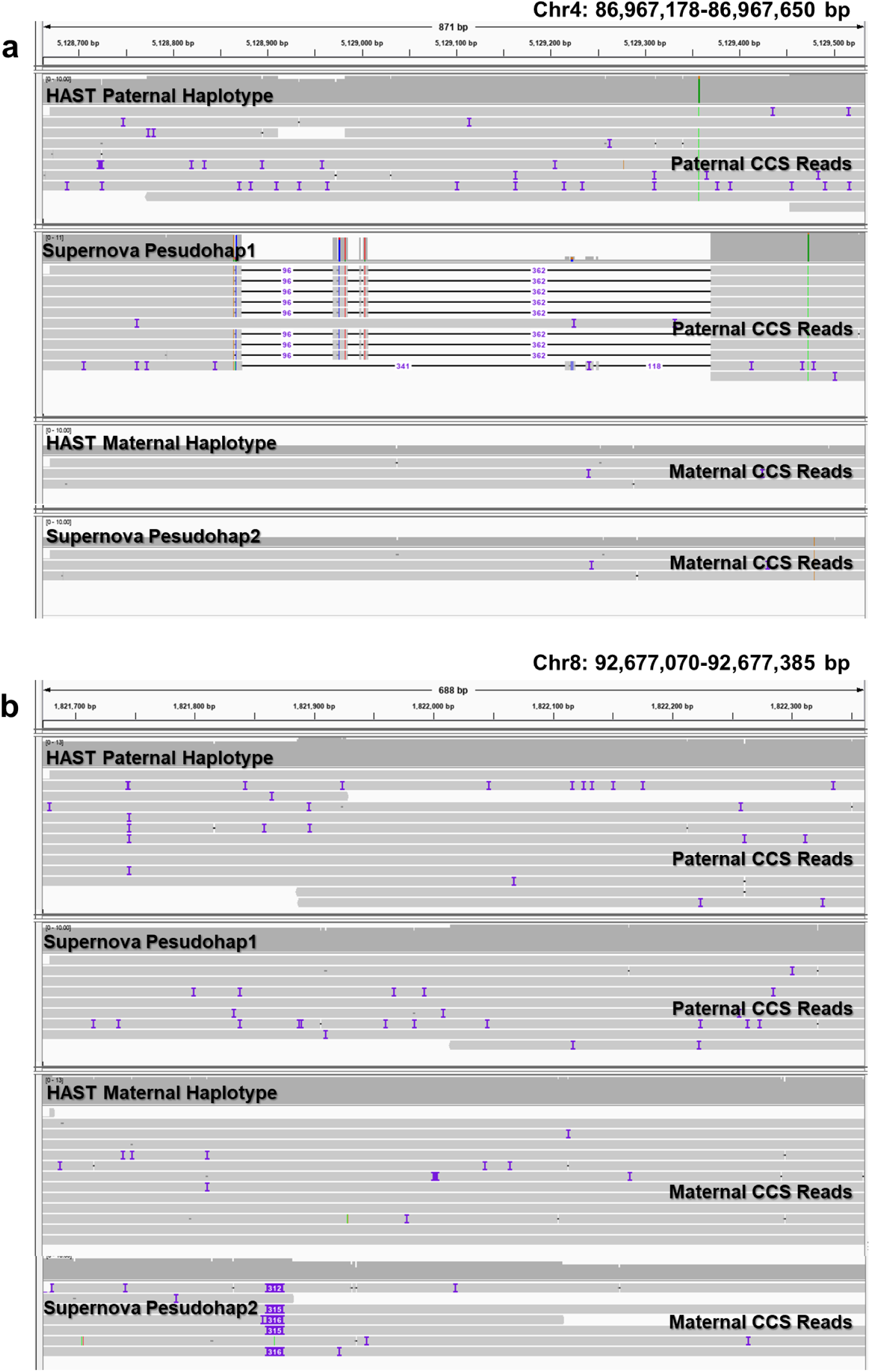
Structural variants visualized in IGV revealed for HAST’s haplotypes and Supernova’s pseudo-haplotypes. The mapped CCS reads with each assembly in the Integrative Genomics Viewer (IGV)^45^ display SV in phase with single-nucleotide variants. The pseudo-haplotypes are missing two phased SVs: **a**) a paternal-specific deletion in hg19 chr4: 86,967,178-86,967,650, and **b**) a maternal-specific insertion in hg19 chr8: 92,677,070-92,677,385. The CCS reads were firstly partitioned by TrioCanu, and then individually mapped to the corresponding assemblies with minimap2^46^. All the assemblies were aligned with the reference to obtain the relation of scaffolds/contigs with chromosomes. We arbitrarily selected several known SV in standard calls, and focused on the segments in the same region.

TrioCanu^12^ also provided two haplotype-resolved assemblies using the trio-binning strategy, with the contig N50 of 726 kb and 1.49 Mb for paternal and maternal, respectively (**Supplementary Table 5**). With mapping against the reference genome, the contig NGA50 and genome coverage ranged from 444 kb to 758 kb, and 90.87% to 96.22%. The variations between paternal and maternal haplotypes reflected the assembly complexity for Y chromosome, which were consistent with HAST. With the benefit of long read length, CCS assemblies exhibited similar haplotyping precision (99.90% and 99.91%) and recall rates (96.12% and 92.78%) relative to HAST, as well as the completeness in BUSCO assessment (88.8% and 92.5%, **Table 1**). Almost all the haplotype-resolved contigs were aligned along either the x-axis or y-axis in the blob plot, which demonstrated the excellent haplotyping accuracy and efficiency with trio binning (**Fig. 3i**). In general, the reconstruction of two haplotypes by TrioCanu was equivalent to that by HAST, and superior to Supernova without trio binning.

Whereas, the average *k*-mer-level accuracy in the CCS assemblies was only Q38 (**Table 1**), obviously lower than stLFR assemblies by HAST or Supernova (>Q60). In spite of the pre-error correction by the consensus procedure, the residuals still introduced a remarkable number of misassembled *k*-mers shown in the copy-number or haplotyping assembly spectrum, with a peak higher than stLFR assemblies (**Fig. 3c** and **3f**). In addition, the recall number of heterozygous *k*-mers for each parent was not as balanced as HAST’s, implying the comparatively larger variations in the assembly quality (**Fig. 3f**). Nevertheless, haplotype-resolved CCS assemblies produced an 18-fold longer contig N50 compared with HAST’s (**Table 1**) but a 10-fold shorter phase block N50 (**Table 2**) in absence of long-range scaffolding information, which implied a possible combination with HAST’s haplotypes.

## Discussion

In this study, we provided a direct way to individually assemble the haplotypes using trio binning. Although the read intensity or spanning length varies for different SLR datasets, HAST clusters reads sharing the same barcodes and retains the long-range phased sequence information. The application of trio binning by HAST simplifies the assembly problems to some extent, and achieves significant improvements in the continuity, accuracy and completeness of haplotyping, but not limited to the diploid genomes. The completeness of sex chromosomes could be further enhanced by the identified sex-specific *k*-mers^47^. It also applies to complex polyploid genomes if the parents or related species are available, thus possibly promoting the animal and plant genetics and breeding. In addition, the true recovery of hereditary information and the accurate ancestor assignment of chromosomes are also essential for studies of structure variations and evolutionary relation.

HAST generates two haplotype-resolved assemblies, but the partitioned long fragments can be used separately. It demonstrates a possibility of combining SLR data with other data type, for instance, PacBio and ONT long reads, as well as other bioinformatics tools, for instance, single-nucleotide-polymorphism or structural-variation callers. A typical association with long reads relates to the extension of scaffolds^48^ and contigs^49, 50^. It is necessary to take full advantages of different sequencing platforms and comprehensively utilize genomic information in short range (short reads, <1kb), medium range (long reads, 1k∼10kb) and long range (ultra-long reads, SLR, Hi-C, BioNano, 10kb∼Mb) with various error rates and balance their costs.

## Methods

### Sample preparation and sequencing of the HJ trios

We sequenced a Han Chinese volunteer (Research ethics ID: XHEC-C-2019-086, HJ) using MGIEasy stLFR Library Prep Kit on MGISEQ-2000 platform with the data size of 632 Gb. The application of SOAPfilter (version 2.2)^51^ removed possible adapter contaminations, low quality reads, duplications, and PE reads with improper insert size, reducing the data size to 355 Gb totally. The haplotype *k*-mers were extracted from his parental MPS short reads, including 29.6× depth of paternal and 29.8× depth of maternal, respectively. Moreover, to construct the haplotypes and compare with HAST’s results, PacBio SMRT libraries were sequenced on a Sequel instrument, which output 870 Gb subreads, and finally generated 66 Gb CCS reads.

### Generation of parental unique markers

The classification of parental unshared *k*-mers is similar to the method described in TrioCanu^12^. Jellyfish^52^(version 0.6.1) was chosen to generate, count, and output distinct *k*-mers in the parental genomes due to its high speed and low requirement of memory. The identification of haplotype *k*-mers was straightforward by mixing 1 copy of paternal *k*-mers and 2 copies of maternal and counting the total frequency. Then the *k*-mers that occurred exactly once in the mixture belonged to the paternal group, while those that occurred twice were maternal-specific.

However, the original number of unshared *k*-mers was surplus, and the majority originated from the sequencing errors (low coverage). The rest of *k*-mers including those with repeat-induced high coverage exclusively represented the parental characteristics in all likelihood. To remove the suspicious unshared *k*-mers and reduce the computational amount, we limited the parental *k*-mer libraries according to the coverage distribution. For the plot of *k*-mer frequency *f* against coverage *c*, the low coverage threshold *C*_*low*_ was determined as the coverage value *C*_*1*_ corresponding to the first lowest point of the curve. The rest of the curve excluding sequencing errors approximately resembled a positive skew Gaussian distribution with a long tail for high coverages, and the degree of dispersion was different in low- and high-coverage regions. For convenience, we assumed the dispersion degree for high coverages was twice of that for low coverages. Then the high coverage threshold *C*_*high*_ was defined by the following formula

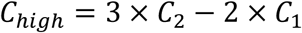

where the coverage *C*_*2*_ corresponded to the first highest point of the curve. Only those *k*-mers within the thresholds were exported to identify long fragments in the next step. Note that the number of unshared *k*-mers is not necessary to be equal for each haplotype due to read errors.

The performance of the classification also substantially depends on the *k* value. Theoretically, a read in LFR has a greater possibility of matching heterozygous markers by smaller *k*-mers. However, a too small *k* value might cause random *k*-mer collisions, especially for large genomes. According to the formula of *k*-mer collision rate *r* given a random distribution of *k*-mers in the genome *G*^53^,

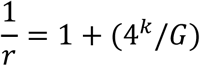

a *k* of 19 provided a collision rate of 1% for the human genome, while *k*=21,31 reduced *r* below 0.1% and 1e^-9^, respectively.

### Partitioning of stLFR long fragments

The concept of read binning prior to assembly has been used in long reads^12^ and metagenomics^54-56^. Ideally, an LFR can be categorized if it possesses individual *k*-mers from one haplotype and none from others. Assuming that heterozygous sites and sequencing errors are randomly distributed in the genome and there are no overlaps between inter- or intra-read pairs in the same LFR, the possibility *p* of the occurrence of one unshared *k*-mer in one LFR can be derived as

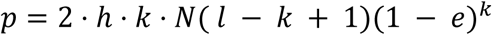

where *h* is the genome heterozygosity, *N* is the number of read pairs in one LFR, *l* is the read length, and *e* is the sequencing error rate. The read clustering based on barcodes overcomes the limit of short read length, especially for genomes with low heterozygosity. The high base-calling accuracy also improves the sensitivity and precision of *k*-mer mapping. Given *h*=0.1%, *N*=20, *l*=100, *k*=21, *e*=0.1% for a typical stLFR library of human, the expected number of unshared *k*-mers is larger than 65 for a barcode.

In the case of trios, LFR’s are assigned to the paternal group if they only own paternal-specific *k*-mers and vice versa. If both types of parental *k*-mers match portions of the same LFR, then it will be judged by the normalized opportunity of unshared *k*-mer mapping, with a correction factor. The factor attempts to neutralize the inherent discrepancy of parents due to the constitution of sex chromosomes^57^. The remaining LFR’s with equal *k*-mer counts are discarded as errors, while those with no *k*-mers are regarded as homozygous. Note that each LFR is identified by its barcode, which corresponds to all fragments in the same co-barcoding compartment^10^. To accelerate millions of *k*-mer queries in the partitioning, we binarized the *k*-mer characters, hashed the *k*-mer database, and parallelized the classifying procedure. Note that HAST also accepts 10X Genomics reads, PacBio and ONT long reads, and shows significant improvements in speed and memory efficiency relative to the trio-binning scripts in the published TrioCanu^12^ (**Supplementary Table 8**). We notice that the latest version of Canu^58^ has also optimized its thread and memory consumptions.

### Haplotyped-resolved assembly pipeline

All LFR’s in paternal or maternal group were combined with homozygous LFR’s, transformed into 10X Genomics format, and passed to Supernova (version 2.1.1)^9^ to assemble, respectively. Supernova utilizes barcode and PE information to solve the repeat-induced regions in the assembly graph, while the heterozygosity-induced problems are forestalled by the trio-binning strategy. Note that the direct pseudo-haplotype output still retains the bubble structures in the graph, and arbitrarily selects one arm for each bubble. We used the haplotyped-specific *k*-mers again to determine which arm truly represented the allelic variants, reported in the final haplotype assembly.

### Haplotyped-resolved assembly for PacBio HiFi CCS data

We used the TrioCanu module implemented in Canu (version 1.9) to classify the PacBio CCS long reads with an average read length of 11.4 kb. The identified ratio was 35.9% and 36.4% for paternal and maternal, respectively (**Supplementary Table 4**). The haplotypes were individually assembled with haplotype-specific reads and those could not be assigned.

### Validation of haplotyping effects

Hg19 assembly^59^ was used as the reference for *H. sapiens*. QUAST^60^ (version 5.0.2) was used with default parameters to report the assembly statistics including total length, scaffold N50, contig N50, as well as scaffold NGA50, contig NGA50, genome fraction, misassemblies and local misassemblies based on the mapping relation with the reference. Note that QUAST split the scaffold by continuous fragments of N’s of length ≥ 10 as contig. In addition, Merqury^44^ (version 1.0) was applied for the evaluation of haplotyping accuracy and completeness according to the reliable specific *k*-mers from the offspring’s and parental reads. In total, we generated 18.6 Mb paternal hap-mers and 16.3 Mb maternal hap-mers to calculate the haplotyping precision and recall. To quantify the effect of HAST on the downstream bioinformatics analysis, we ran BUSCO^61^ (version 3.0.2) analysis for all the human assemblies against the vertebrata_odb9.

## Author contributions

XXX

## Data availability

XXX

## Code availability

All codes and scripts to build haplotype-specific *k*-mer sets, and classify stLFR reads are available at https://github.com/BGI-Qingdao/HAST.

## Authors’ contributions

XXX

## Supplementary Material

XXX

## Acknowledgements

XXX

## References

1. Tewhey, R., Bansal, V., Torkamani, A., Topol, E.J. & Schork, N.J. The importance of phase information for human genomics. Nat Rev Genet 12, 215–223 (2011).

2. Muers, M. Genomics: No half measures for haplotypes. Nat Rev Genet 12, 77 (2011).

3. Yang, J. et al. Haplotype-resolved sweet potato genome traces back its hexaploidization history. Nat Plants 3, 696–703 (2017).

4. Myers, E.W. et al. A whole-genome assembly of Drosophila. Science 287, 2196–2204 (2000).

5. Lander, E.S. et al. Initial sequencing and analysis of the human genome. Nature 409, 860–921 (2001).

6. Yang, H., Chen, X. & Wong, W.H. Completely phased genome sequencing through chromosome sorting. Proc Natl Acad Sci U S A 108, 12–17 (2011).

7. Peters, B.A. et al. Accurate whole-genome sequencing and haplotyping from 10 to 20 human cells. Nature 487, 190–195 (2012).

8. Zheng, G.X. et al. Haplotyping germline and cancer genomes with high-throughput linked-read sequencing. Nat Biotechnol 34, 303–311 (2016).

9. Weisenfeld, N.I., Kumar, V., Shah, P., Church, D.M. & Jaffe, D.B. Direct determination of diploid genome sequences. Genome Res 27, 757–767 (2017).

10. Wang, O. et al. Efficient and unique cobarcoding of second-generation sequencing reads from long DNA molecules enabling cost-effective and accurate sequencing, haplotyping, and de novo assembly. Genome Res 29, 798–808 (2019).

11. Chin, C.S. et al. Phased diploid genome assembly with single-molecule real-time sequencing. Nat Methods 13, 1050–1054 (2016).

12. Koren, S. et al. De novo assembly of haplotype-resolved genomes with trio binning. Nat Biotechnol (2018).

13. Selvaraj, S., J, R.D., Bansal, V. & Ren, B. Whole-genome haplotype reconstruction using proximity-ligation and shotgun sequencing. Nat Biotechnol 31, 1111–1118 (2013).

14. Zhang, X., Zhang, S., Zhao, Q., Ming, R. & Tang, H. Assembly of allele-aware, chromosomal-scale autopolyploid genomes based on Hi-C data. Nat Plants 5, 833–845 (2019).

15. Seo, J.S. et al. De novo assembly and phasing of a Korean human genome. Nature 538, 243–247 (2016).

16. Patterson, M. et al. WhatsHap: Weighted Haplotype Assembly for Future-Generation Sequencing Reads. J Comput Biol 22, 498–509 (2015).

17. Edge, P., Bafna, V. & Bansal, V. HapCUT2: robust and accurate haplotype assembly for diverse sequencing technologies. Genome Res 27, 801–812 (2017).

18. Kuleshov, V. et al. Whole-genome haplotyping using long reads and statistical methods. Nat Biotechnol 32, 261–266 (2014).

19. Kim, D., Paggi, J.M., Park, C., Bennett, C. & Salzberg, S.L. Graph-based genome alignment and genotyping with HISAT2 and HISAT-genotype. Nat Biotechnol 37, 907–915 (2019).

20. Bishara, A. et al. Read clouds uncover variation in complex regions of the human genome. Genome Res 25, 1570–1580 (2015).

21. Zhang, X., Wu, R., Wang, Y., Yu, J. & Tang, H. Unzipping haplotypes in diploid and polyploid genomes. Comput Struct Biotechnol J 18, 66–72 (2020).

22. Kronenberg, Z.N. et al. Extended haplotype phasing of <em>de novo</em> genome assemblies with FALCON-Phase. bioRxiv, 327064 (2019).

23. Garg, S. et al. Efficient chromosome-scale haplotype-resolved assembly of human genomes. bioRxiv, 810341 (2019).

24. Low, W.Y. et al. Haplotype-resolved genomes provide insights into structural variation and gene content in Angus and Brahman cattle. Nat Commun 11, 2071 (2020).

25. Marchini, J. et al. A comparison of phasing algorithms for trios and unrelated individuals. Am J Hum Genet 78, 437–450 (2006).

26. Eberle, M.A. et al. A reference data set of 5.4 million phased human variants validated by genetic inheritance from sequencing a three-generation 17-member pedigree. Genome Res 27, 157–164 (2017).

27. Campoy, J.A. et al. Chromosome-level and haplotype-resolved genome assembly enabled by high-throughput single-cell sequencing of gamete genomes. bioRxiv, 2020.2004.2024.060046 (2020).

28. Low, W.Y. et al. Haplotype-resolved genomes provide insights into structural variation and gene content in Angus and Brahman cattle. Nature Communications 11, 2071 (2020).

29. Peters, B.A. et al. Accurate whole-genome sequencing and haplotyping from 10 to 20 human cells. Nature 487, 190 (2012).

30. Amini, S. et al. Haplotype-resolved whole-genome sequencing by contiguity-preserving transposition and combinatorial indexing. Nature genetics 46, 1343 (2014).

31. Zheng, G.X. et al. Haplotyping germline and cancer genomes with high-throughput linked-read sequencing. Nature biotechnology 34, 303 (2016).

32. Kuleshov, V. et al. Whole-genome haplotyping using long reads and statistical methods. Nature biotechnology 32, 261 (2014).

33. Marks, P. et al. Resolving the full spectrum of human genome variation using Linked-Reads. Genome research (2019).

34. Bishara, A. et al. Read clouds uncover variation in complex regions of the human genome. Genome research 25, 1570–1580 (2015).

35. Elyanow, R., Wu, H.-T. & Raphael, B.J. Identifying structural variants using linked-read sequencing data. Bioinformatics 34, 353–360 (2017).

36. Zhou, X. et al. Aquila stLFR: assembly based variant calling package for stLFR and hybrid assembly for linked-reads. bioRxiv, 742239 (2019).

37. Adey, A. et al. In vitro, long-range sequence information for de novo genome assembly via transposase contiguity. Genome research 24, 2041–2049 (2014).

38. Kuleshov, V., Snyder, M.P. & Batzoglou, S. Genome assembly from synthetic long read clouds. Bioinformatics 32, i216–i224 (2016).

39. Weisenfeld, N.I., Kumar, V., Shah, P., Church, D.M. & Jaffe, D.B. Direct determination of diploid genome sequences. Genome research 27, 757–767 (2017).

40. Yeo, S., Coombe, L., Warren, R.L., Chu, J. & Birol, I. ARCS: scaffolding genome drafts with linked reads. Bioinformatics 34, 725–731 (2017).

41. Coombe, L. et al. ARKS: chromosome-scale scaffolding of human genome drafts with linked read kmers. BMC bioinformatics 19, 234 (2018).

42. Fan, G. et al. Initial data release and announcement of the Fish10K: Fish 10,000 Genomes Project. bioRxiv, 787028 (2019).

43. Xu, M. et al. Draft genome of a porcupinefish, Diodon Holocanthus. bioRxiv, 775387 (2019).

44. Rhie, A., Walenz, B.P., Koren, S. & Phillippy, A.M. Merqury: reference-free quality and phasing assessment for genome assemblies. bioRxiv, 2020.2003.2015.992941 (2020).

45. Robinson, J.T. et al. Integrative genomics viewer. Nat Biotechnol 29, 24–26 (2011).

46. Li, H. Minimap2: pairwise alignment for nucleotide sequences. Bioinformatics 34, 3094–3100 (2018).

47. Wang, X., Liu, Q., Li, A. & Ruan, J. SRY: An Effective Method for Sorting Long Reads of Sex-limited Chromosome. bioRxiv, 2020.2005.2025.115592 (2020).

48. Boetzer, M. & Pirovano, W. SSPACE-LongRead: scaffolding bacterial draft genomes using long read sequence information. BMC Bioinformatics 15, 211 (2014).

49. Xu, M. et al. TGS-GapCloser: fast and accurately passing through the Bermuda in large genome using error-prone third-generation long reads. bioRxiv, 831248 (2019).

50. Murigneux, V. et al. Comparison of long read methods for sequencing and assembly of a plant genome. bioRxiv, 2020.2003.2016.992933 (2020).

51. Luo, R. et al. SOAPdenovo2: an empirically improved memory-efficient short-read de novo assembler. Gigascience 1, 18 (2012).

52. Marcais, G. & Kingsford, C. A fast, lock-free approach for efficient parallel counting of occurrences of k-mers. Bioinformatics 27, 764–770 (2011).

53. Fofanov, Y. et al. How independent are the appearances of n-mers in different genomes? Bioinformatics 20, 2421–2428 (2004).

54. Wu, Y.W. & Ye, Y. A novel abundance-based algorithm for binning metagenomic sequences using l-tuples. J Comput Biol 18, 523–534 (2011).

55. Wang, Y., Leung, H.C., Yiu, S.M. & Chin, F.Y. MetaCluster 5.0: a two-round binning approach for metagenomic data for low-abundance species in a noisy sample. Bioinformatics 28, i356–i362 (2012).

56. Cleary, B. et al. Detection of low-abundance bacterial strains in metagenomic datasets by eigengenome partitioning. Nat Biotechnol 33, 1053–1060 (2015).

57. Bachtrog, D. & Charlesworth, B. Towards a complete sequence of the human Y chromosome. Genome Biol 2, Reviews1016 (2001).

58. Koren, S. et al. Canu: scalable and accurate long-read assembly via adaptive k-mer weighting and repeat separation. Genome Res 27, 722–736 (2017).

59. Church, D.M. et al. Modernizing reference genome assemblies. PLoS Biol 9, e1001091 (2011).

60. Gurevich, A., Saveliev, V., Vyahhi, N. & Tesler, G. QUAST: quality assessment tool for genome assemblies. Bioinformatics 29, 1072–1075 (2013).

61. Simao, F.A., Waterhouse, R.M., Ioannidis, P., Kriventseva, E.V. & Zdobnov, E.M. BUSCO: assessing genome assembly and annotation completeness with single-copy orthologs. Bioinformatics 31, 3210–3212 (2015).

