## Supplemental Information for "Haplotype-Resolved Assembly for Synthetic Long Reads Using a Trio-Binning Strategy"

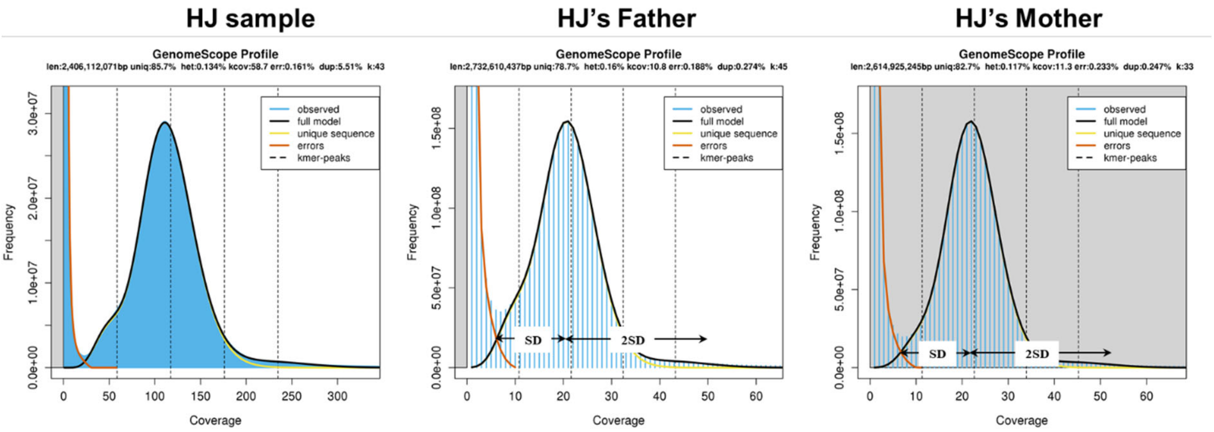


**Supplementary Figure 1.** The *k*-mer coverage histogram graphs for the offspring and his parents, and the schematic of cutoff coverages for the generation of haplotype-specific *k*-mer libraries.

**Supplementary Table 1.** Haplotype-specific $k$-mers with different $k$-values. The frequency range is given by the $k$-mer frequency histogram. The upper limit is calculated by the formula $Upper\ Limit = k \cdot h \cdot G$ where $h$ refers to the genome heterozygosity, and $G$ is the genome size.

| | **21-mers** | | | **19-mers** | | | **31-mers** | | |
|---|---|---|---|---|---|---|---|---|---|
| | **Frequency Range** | **Haplotype-specific** | **Upper Limit** | **Frequency Range** | **Haplotype-specific** | **Upper Limit** | **Frequency Range** | **Haplotype-specific** | **Upper Limit** |
| **Paternal** | 9-58 | 62,001,448 | 63M | 10-54 | 53,587,552 | 57M | 9-52 | 83,254,249 | 93M |
| **Maternal** | 9-58 | 39,206,623 | 63M | 10-54 | 31,661,160 | 57M | 9-52 | 66,808,818 | 93M |

**Supplementary Table 2.** The partitioned ratio of total barcodes and reads after three iterations using different *k*-values. Paternal: only have paternal-specific *k*-mers; Maternal: only have maternal-specific *k*-mers; Shared: have both types of parent-specific *k*-mers; Homozygous: have no parental-specific *k*-mers.

| | **21-mer** | **19-mer** | **31-mer** | **Final** |
|---|---|---|---|---|
| **Haploid Barcodes** | | | | |
| Paternal | 9.2% | 2.3% | 0.8% | 12.2% |
| Maternal | 9.2% | 1.3% | 1.0% | 11.6% |
| Shared | 2.3% | 0.3% | 0.0% | 2.7% |
| Homozygous | 79.3% | 75.3% | 73.5% | 73.5% |
| **Haploid Reads Clustered by Barcodes** | | | | |
| Paternal | 18.9% | 6.1% | 1.2% | 26.2% |
| Maternal | 19.3% | 3.6% | 1.7% | 24.6% |
| Shared | 11.5% | 1.5% | 0.1% | 13.1% |
| Homozygous | 42.3% | 38.0% | 36.1% | 36.1% |
| **Haploid Reads without Barcodes** | | | | |
| Paternal | 1.5% | - | - | - |
| Maternal | 1.4% | - | - | - |
| Shared | 0.0% | - | - | - |
| Homozygous | 97.1% | - | - | - |

**Supplementary Table 3.** The mapping statistics for classified stLFR reads against the reference genome and TrioCanu-assembled haplotypes. Paternal: only have paternal-specific *k*-mers; Maternal: only have maternal-specific *k*-mers; Shared: have both types of parent-specific *k*-mers; Homozygous: have no parent-specific *k*-mers. All results were reported by stLFRQC (https://github.com/BGI-Qingdao/stLFRQC).

| | **Paternal** | **Maternal** | **Shared** | **Homozygous** | **Paternal+ Homozygous** | **Maternal+ Homozygous** |
|---|---|---|---|---|---|---|
| **Reference: hg19** | | | | | | |
| Sequencing Depth (X) | 26.3 | 27.8 | 14.6 | 26.6 | 52.9 | 54.4 |
| Genome Coverage (%) | 41.4 | 46.1 | 3.1 | 41.6 | 83.0 | 87.7 |
| Mismatch (%) | 0.72 | 0.71 | 0.73 | 0.83 | - | - |
| Indel (%) | 0.06 | 0.06 | 0.05 | 0.05 | - | - |
| **Reference: Paternal Assembly by Pacbio CCS** | | | | | | |
| Sequencing Depth (X) | 30.5 | 31.7 | 16.8 | 31.0 | 61.5 | 62.7 |
| Genome Coverage (%) | 49.5 | 53.7 | 5.2 | 49.1 | 98.6 | 100.0 |
| Mismatch (%) | 0.49 | 0.62 | 0.56 | 0.65 | - | - |
| Indel (%) | 0.03 | 0.06 | 0.05 | 0.04 | - | - |
| **Reference: Maternal Assembly by Pacbio CCS** | | | | | | |
| Sequencing Depth (X) | 28.8 | 30.7 | 16.1 | 29.8 | 58.6 | 60.5 |
| Genome Coverage (%) | 45.7 | 51.3 | 4.40 | 46.3 | 92.0 | 97.6 |
| Mismatch (%) | 0.60 | 0.46 | 0.52 | 0.61 | - | - |
| Indel (%) | 0.06 | 0.03 | 0.02 | 0.03 | - | - |

**Supplementary Table 4.** The LFR collision rate with different reference assemblies. Paternal: only have paternal-specific *k*-mers; Maternal: only have maternal-specific *k*-mers; Shared: have both types of parent-specific *k*-mers; Homozygous: have no parent-specific *k*-mers. Read pairs with the same barcode occurring in the same 300 kb range of reference genome are considered from one LFR. LFR's have less than 5 read pairs were filtered in the calculation. The results were reported by stLFRQC (https://github.com/BGI-Qingdao/stLFRQC).

| | **Paternal** | **Maternal** | **Shared** | **Homozygous** | **Paternal+ Homozygous** | **Maternal+ Homozygous** |
|---|---|---|---|---|---|---|
| **Reference: hg19** | | | | | | |
| LFR Collision Rate | 1.58 | 1.54 | 2.12 | 1.19 | 1.38 | 1.37 |
| **Reference: Paternal Assembly by Pacbio CCS** | | | | | | |
| LFR Collision Rate | 1.72 | 1.80 | 2.46 | 1.27 | 1.49 | 1.54 |
| **Reference: Maternal Assembly by Pacbio CCS** | | | | | | |
| LFR Collision Rate | 1.67 | 1.71 | 2.35 | 1.24 | 1.45 | 1.48 |

**Supplementary Table 5.** The statistics for haplotype-resolved Pacbio CCS reads. CCS long reads were partitioned by the trio-binning module in Canu (version 1.9).

| | Total Length (bp) | Total Reads (#) | Average Length (bp) | N50 Length (bp) | Maximum Length (bp) | Minimum Length (bp) |
|---|---|---|---|---|---|---|
| **Paternal** | 16,725,028,348 | 1,399,172 | 11,954 | 12,827 | 21,532 | 1,000 |
| **Maternal** | 16,944,352,807 | 1,412,009 | 12,000 | 12,897 | 21,824 | 1,000 |
| **Unknown** | 12,888,057,009 | 1,248,031 | 10,327 | 11,042 | 21,116 | 1,000 |

**Supplementary Table 6.** The LFR statistics for HAST and Supernova.

| | **Paternal** | **Maternal** | **Pseudohap1** | **Pseudohap2** |
|---|---|---|---|---|
| **Assembler** | HAST | HAST | Supernova | Supernova |
| **Raw COV (×)** | 79.35 | 80.58 | 84.55 | 84.55 |
| **EFFECTIVE COV (×)** | 47.36 | 48.19 | 49.72 | 49.72 |
| **MOLECULE LEN (kbp)** | 118.25 | 119.43 | 128.04 | 128.04 |
| **BARCODE N50 (read)** | 116 | 122 | 556 | 556 |
| **DUPS (%)** | 31.41 | 31.28 | 32.53 | 32.53 |

**Supplementary Table 7.** The haplotype-resolved assembly statistics for HAST, and comparisons to the direct Supernova outputs and TrioCanu assemblies. HAST version (1.0.0) was used with a modified barcode list. The version 2.1.1 of Supernova was used for pseudo-haplotypes. Canu (version 1.9) was used to generate two haplotypes. All assemblies were run with default parameters.

| | **Paternal** | **Maternal** | **Pseudohap1** | **Pseudohap2** | **Paternal** | **Maternal** |
|---|---|---|---|---|---|---|
| **Assembler** | HAST | HAST | Supernova | Supernova | TrioCanu | TrioCanu |
| **Total Length (bp)** | 3,007,368,641 | 3,000,964,520 | 3,163,445,831 | 3,163,254,706 | 2,756,168,090 | 2,922,794,873 |
| **Genome Fraction (%)** | 94.392 | 94.947 | 94.278 | 94.278 | 90.871 | 96.224 |
| **Scaffold N50 (bp)** | 11,419,520 | 18,265,876 | 16,663,863 | 16,660,859 | - | - |
| **Scaffold NGA50 (bp)** | 1,019,668 | 1,141,819 | 954,188 | 948,377 | - | - |
| **Contig N50 (bp)** | 60,970 | 63,759 | 37,562 | 37,553 | 726,347 | 1,489,551 |
| **Contig NGA50 (bp)** | 50,989 | 54,014 | 31,642 | 31,658 | 444,469 | 758,832 |
| **# misassemblies** | 6,260 | 5,861 | 6,231 | 6,199 | 5,663 | 6,284 |
| **# local misassemblies** | 7,300 | 6,142 | 6,096 | 6,117 | 8,996 | 9,427 |
| **# mismatches /100 kbp** | 115 | 114 | 112 | 112 | 135 | 136 |
| **# indels /100 kbp** | 26 | 26 | 25 | 25 | 52 | 46 |

**Supplementary Table 8.** Runtime and memory requirement by HAST (HAST4TGS) and TrioCanu for the human Pacbio reads. All tests were run with 30 threads on the same machine. In total, 30× MPS reads for each parent were used to build the unshared *k*-mer libraries, with which about 22× Pacbio CCS long reads were partitioned. The subversion of HAST (HAST4TGS, https://github.com/BGI-Qingdao/HAST4TGS) was used to support long reads. TrioCanu was run according to the example scripts from https://github.com/skoren/triobinningScripts.

| | | **HAST** | **TrioCanu** |
|---|---|---|---|
| | **Wall-clock time** | 13.5 h | 18.8 h |
| *Generate k-mers* | **CPU time** | 47.4 h | 168.0 h |
| | **Memory** | 51 GB | 1,669 GB |
| | **Wall-clock time** | 1.5 h | 236.4 h |
| *Classify reads* | **CPU time** | 36.4 h | 236.4 h |
| | **Memory** | 22 GB | 91 GB |
| | **Wall-clock time** | 15.0 h | 255.2 h |
| *Total* | **CPU time** | 83.8 h | 404.4 h |
| | **Memory** | 51 GB | 1,669 GB |
